# A subset of sweet-sensing neurons identified by IR56d are necessaryand sufficient for fatty acid taste

**DOI:** 10.1101/174623

**Authors:** John M. Tauber, Elizabeth Brown, Yuanyuan Li, Maria E. Yurgel, Pavel Masek, Alex C. Keene

## Abstract

Fat represents a calorically potent food source that yields approximately twice the amount of energy as carbohydrates or proteins per unit of mass. The highly palatable taste of FAs promotes food consumption, activates reward circuitry, and is thought to contribute to hedonic feeding underlying many metabolism-related disorders. Despite a role in the etiology of metabolic diseases, little is known about how dietary fats are detected by the gustatory system to promote feeding. Previously, we showed that a broad population of sugar-sensitive taste neurons expressing Gustatory Receptor 64f (*Gr64f*), is required for reflexive feeding responses to both FAs and sugars. Here, we report a genetic silencing screen to identify specific populations of taste neurons that mediate fatty acid taste. We find neurons identified by expression of Ionotropic Receptor 56d (*IR56d*) are necessary and sufficient for reflexive feeding response to FAs. Functional imaging reveals that *IR56d*-expressing neurons are responsive to short- and medium-chain FAs. Silencing *IR56d* neurons selectively abolishes fatty acid taste, while their activation is sufficient to drive feeding responses. Analysis of co-expression with *Gr64f* identifies two subpopulations of *IR56d*-expressing neurons. While physiological imaging reveals that both populations are responsive to FAs, *IR56d*/*Gr64f* neurons are activated by medium-chain FAs and are sufficient for reflexive feeding responses to FA. Moreover, flies can discriminate between sugar and FAs in an aversive taste memory assay, indicating that FA taste is a unique modality in *Drosophila*. Taken together, these findings localize fatty acid taste within the *Drosophila* gustatory center and provide an opportunity to investigate discrimination between different categories of appetitive tastants.

**Authors Summary:** Fat represents a calorically potent food source that yields approximately twice the amount of energy as carbohydrates or proteins per unit of mass. Dietary lipids are comprised of both triacylglycerides and FAs, and growing evidence suggests that it is the free FAs that are detected by the gustatory system. The highly palatable taste of FAs promotes food consumption, activates reward centers in mammals, and is thought to contribute to hedonic feeding that underlies many metabolism-related disorders. Despite a role in the etiology of metabolic diseases, little is known about how dietary fats are detected by the gustatory system to promote feeding. We have identified a subset of sugar-sensing neurons in the fly that also respond to medium-chain FAs and are necessary and sufficient for behavioral responses to FAs. Further, we find that despite being sensed by shared neuron populations, flies can differentiate between the taste of sugar and FAs, fortifying the notion that FAs and sugar represent distinct taste modalities in flies.

## Introduction

Fat represents a calorically potent food source that yields approximately twice the amount of energy as carbohydrates or proteins per unit of mass. In mammals, dietary lipids are detected by taste cells, mechanosensory and olfactory neurons, as well as by post-ingestive feedback [1–4]. Dietary lipids are comprised of triacylglycerides and FAs (FAs), and growing evidence suggests that it is the free FAs that are detected by the gustatory system [5–7]. Sensing of FAs promotes food consumption, activates reward circuitry, and is thought to contribute to hedonic feeding that underlies many metabolism-related disorders [8,9]. Despite a role in the etiology of metabolic diseases, little is known about how dietary fats are detected by the gustatory system to promote feeding.

In flies and mammals, tastants are sensed by dedicated gustatory receptors that localize to the taste cells or taste receptor neurons [10–12]. These cells are sensitive to different taste modalities such as sweet, bitter, salty, sour or umami, and project to higher order brain structures for processing [10,13,14]. While these taste modalities have been extensively studied, much less is known about how FAs are detected and how information about this sensory stimulus is processed. Taste neurons in *Drosophila* are housed in gustatory sensilla located on the tarsi (feet), proboscis (mouth) and wings. Each sensillum contains dendrites of up to four gustatory receptor neurons (GRNs), which are activated by different categories of tastants [16]. Two main classes of neurons include one group that senses sweet tastants and promotes feeding, and a second, non-overlapping group that senses bitter tastants and promotes food avoidance [17,18]. We have previously shown that *Drosophila* is attracted to medium-chain FAs [19]. Consumption of FAs relies on taste, rather than smell, as it is not impaired by surgical ablation of olfactory organs [19]. Additionally, FA consumption is abolished in *pox-neuro* mutants in which all external taste hairs are converted to mechanosensory bristles, indicating that the chemical signature rather than oily texture of FAs is associated with perception [19]. Silencing *Gr64f* neurons, which are required for sugar sensing [20,21], abolishes behavioral response to FAs, suggesting that shared populations of gustatory neurons detect FAs and sugars [19]. Unlike sugars, FAs sensing is dependent on functional Phospholipase C (PLC) indicating independent sensory pathways for FA and sugar perceptions [18]. However, any further characteristics of the physiological response or the specific neuronal identity of the neurons mediating FA response are unknown.

Taste neurons from the legs and proboscis predominantly project to the subeosophageal zone (SEZ), the primary taste center of the *Drosophila* central nervous system, but the downstream central brain circuitry contributing to taste processing is less well understood [22–25]. Determining how diverse tastants activate GRNs that convey information to the SEZ, and how this information is represented in higher order brain centers, is central to understanding the neural basis for taste processing and feeding behavior. Identifying the neural principles underlying FA taste processing requires identifying the precise populations of taste neurons responsive to FAs, and their innervation of the primary taste center. Recent studies in *Drosophila* have identified taste neurons that are responsive to diverse modalities including salt, sugar, amino acids, water, carbonation, bitter, polyamines, and electrophilic tastants [17,26–32], yet little is known about the populations underlying FA taste. Here, we show that the IR56d population of gustatory neurons, which partially overlaps with Gr64f neurons, is necessary and sufficient to produce a feeding response to FAs. The IR56d-Gr64f population of neurons is selectively responsive to medium-short-chain FAs. Our results reveal a defined channel that senses FAs to promote food consumption, providing a mechanism for differentiation between attractive tastants of different modalities.

## Results

### Hexanoic acid elicits calcium activity in Gr64f sensory neurons

We previously reported that silencing the Gr64f population of primary taste neurons abolishes behavioral response to both sugars and FAs [10]. To investigate the responsiveness of these neurons to FAs directly, the Ca^2+^ sensor GCaMP5 was expressed under control of Gr64f-GAL4 [33,34] (Fig. 1A). The Ca^2+^ response to proboscis application of water, sucrose, or the medium-chain fatty acid hexanoic acid (HxA) was monitored *in vivo* in the axonal projections of Gr64f neurons within the SEZ (Fig. 1B). Flies were provided either 10mM sucrose or 1% HxA, as these concentrations induce comparable levels of proboscis extension reflex behavior. Both 10mM sucrose and 1% HxA induced robust activity in the SEZ, while little response was observed to water alone (Fig. 1C-F). The temporal dynamics of Ca^2+^ activity differed between the two tastants, with HxA eliciting a broader response (Fig. 1E, F), yet both elicited comparable peak changes in fluorescence (Fig 1C). Therefore, both sugars and FAs activate Gr64f sweet-sensing GRNs, fortifying the notion that this neuronal class is generally responsive to attractive tastants.

**Fig 1.**
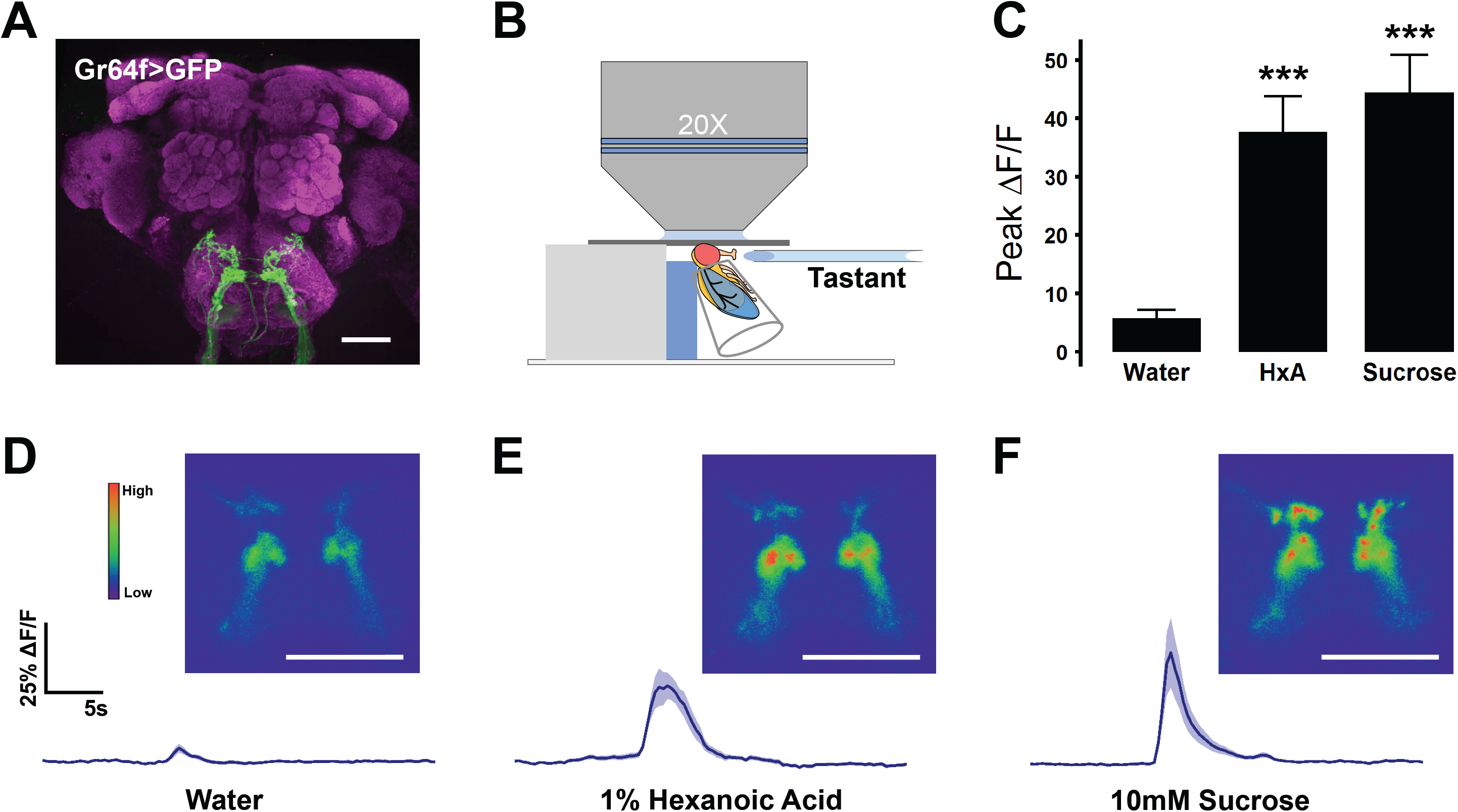
Gr64f gustatory receptor neurons respond to sucrose and HxA. **(A)** Expression of GFP in Gr64f-GAL4 neurons reveals axon terminals in the sub-oesophagael zone (SEZ). Scale bar = 50 μm. **(B)** Diagram of live-imaging experimental protocol. Cuticle above the SEZ is removed and GCaMP5 fluorescence is recorded while tastant is applied to the proboscis. **(C)** Average peak ΔF/F during response to water, 1% HxA, and 10mM sucrose (n = 11, 11, 10 respectively). Error bars indicate SEM. One-way ANOVA with Tukey’s HSD - ***p<0.001 **(D)** Average ΔF/F traces and representative images of calcium activity in Gr64f neurons responding to water, **(E)** 1 % HxA, and **(F)** 10mM sucrose. Scale bar = 50 μm. Shaded region of trace indicates +/- SEM.

To localize FA-sensitive neurons within the broad Gr64f population, we selectively silenced populations of taste neurons predicted to overlap with Gr64f neurons and examined the proboscis extension reflex (PER) in response to sucrose and HxA. The synaptobrevin cleavage peptide Tetanus Toxin-Light Chain (TNT) was expressed under the control of Gustatory Receptor or Ionotropic Receptor promoters known to overlap with Gr64f (Table S1) [20,30,35,36]. Of the 10 GAL4 lines tested, silencing with GAL4 drivers for Gr61a, IR56b, and Gr64f resulted in PER defects to sucrose and HxA, By contrast, silencing Gr64e neurons significantly reduced response to sucrose without affecting response to HxA, and silencing Gr5a, Gr43a, or IR56d neurons significantly reduced PER to HxA without affecting sucrose response.

We chose to further investigate the role of IR56d neurons in FA sensing, since IRs have been found to be involved in detection of non-sweet appetitive tastants, including salt and amino acids [28,37,38]. *IR56d* neurons have previously been reported to project to two distinct regions in the SEZ. One region overlaps with sweet-sensing projections, and the other is indicated to originate in the taste pegs of the proboscis [36]. Outside of sensing carbonation, little is known about ligands that activate the taste pegs or the role they play in the *Drosophila* gustatory system. The sweet-sensing projections of IR56d resembled the region of Gr64f projections that was activated by HxA. Consistent with previous reports, expression of GFP in IR56d neurons (IR56d-GAL4>cd8::GFP) revealed two populations of projections: one innervating the anterior SEZ and previously defined as emanating from labellar bristles, and a second population, emanating from the taste pegs, innervating the posterior SEZ (Fig 2A-C) [36]. To further analyze the role of IR56d neurons, we silenced them using TNT and tested PER upon application of tastants to the proboscis for consistency with live imaging experiments. To control for genetic background and any potential non-specific effects of TNT, we compared PER in flies with silenced IR56d neurons (IR56d-GAL4>UAS-TNT) to flies expressing an inactive variant of TNT (IR56d-GAL4>imp-TNT) [35]. Expression of imp-TNT in IR56d neurons did not affect PER in response to sucrose or HxA, while expression of TNT selectively inhibited HxA response (Fig 2D). Therefore, IR56d neurons are required for behavioral response to HxA, but dispensable for response to sucrose.

**Fig 2.**
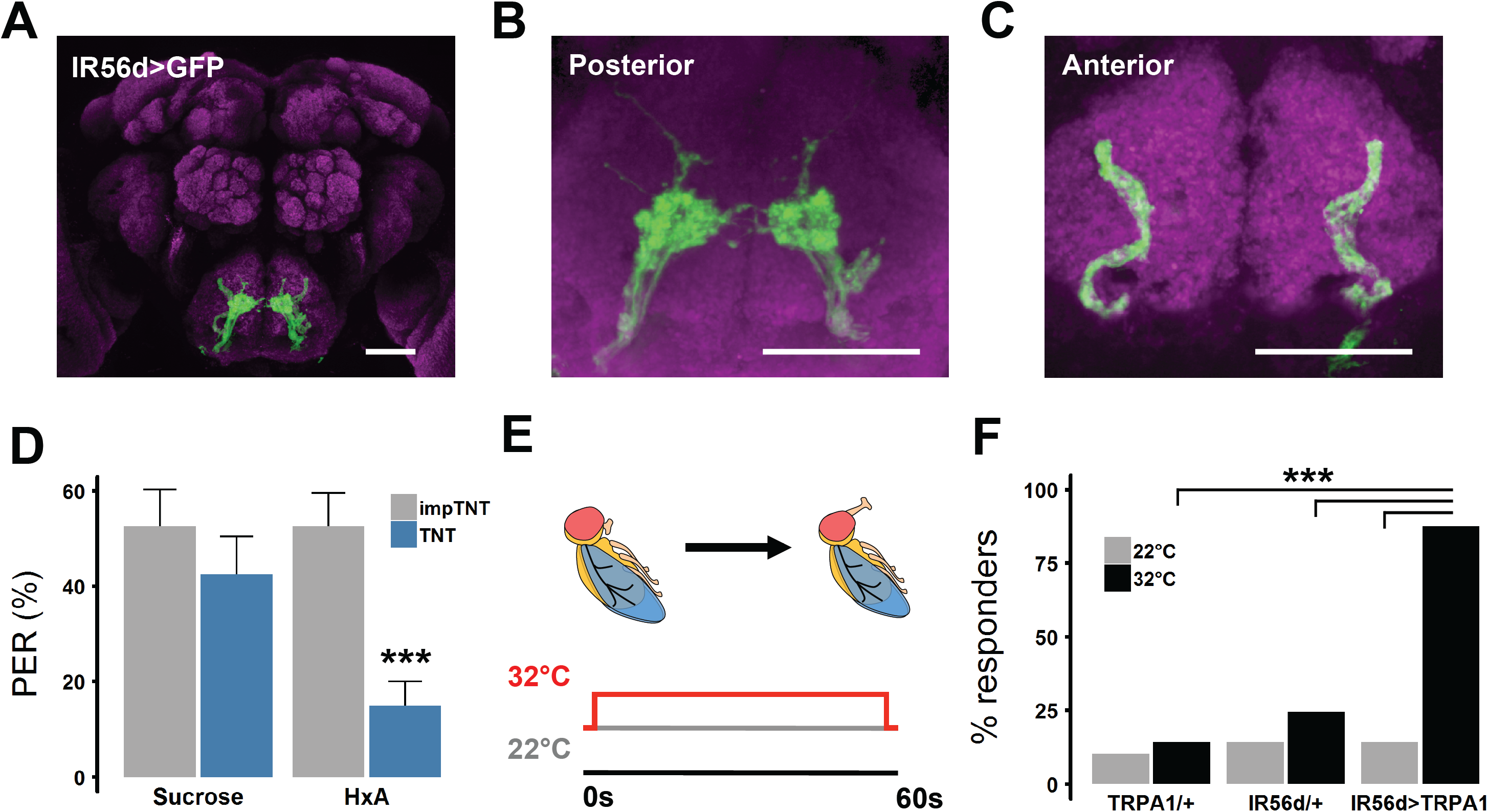
IR56d neurons are necessary and sufficient for PER to FAs. **(A)** Expression pattern of IR56d neurons visualized with GFP. Distinct regions of projection are seen in **(B)** posterior and **(C)** anterior regions of SEZ. Scale bar = 50 μm. **(D)** Blocking synaptic release by genetic expression of light-chain tetanus toxin (TNT) in IR56d neurons significantly reduces PER to HxA, but not sucrose, compared to control flies expressing an inactive form of tetanus toxin (imp-TNT). (imp-TNT n = 26; TNT n = 29). Wilcoxon Rank Sum Test - ***p<0.001. **(E)** Schematic for heat activation of TRPA1. Temperature was elevated for a period of 1 minute and the number of flies showing a PER during this period was counted. **(F)** Heat activation of IR56d neurons with TRPA1 induces significant PER compared to either transgene alone (n = 27 for all genotypes). Fisher’s Exact Test with Bonferroni correction for multiple comparisons - ***p< 0.001

Broad activation of sweet-sensing neurons expressing the trehalose receptor Gr5a induces feeding response in the absence of tastants [17,39,40]. To determine whether activation of IR56d neurons is sufficient to induce PER, we selectively expressed the thermo-sensitive cation channel *transient receptor potential* A1 (TRPA1) in IR56d neurons, or Gr5a neurons as a positive control, and measured heat-induced PER [41,42]. TRPA1 expression induces neuronal activity at temperatures above 28°C, but has little effect on neuronal activity in flies at 22°C, allowing for thermogenetic modulation of neuronal activity [41,42]. In agreement with previous findings, broad activation of sweet-sensing neurons induced robust PER (Fig 2F) [27,43,44]. Similarly, selective activation of IR56d neurons robustly induced PER as compared to control flies harboring *UAS-TRPA1* or *IR56d-GAL4* alone (Fig 2F). Therefore, activation of IR56d neurons alone is sufficient to induce PER.

IR56d neurons project to both taste peg and sweet-sensing regions of the SEZ, and each region can be distinguished anatomically (Fig 2A-C; [36]). To determine whether sugars and FAs can differentially activate the two regions, we expressed GCaMP5 in IR56d neurons (IR56d-GAL4>GCaMP5) and measured tastant-evoked activity separately within the areas of anterior and posterior projections. The anterior region, which overlaps with Gr64f neurons, responded to both HxA and sucrose with similar magnitude (Fig 3A-B). The projections from the taste pegs, however, showed a strong response to HxA, but did not respond to sucrose (Fig 3C-D). Therefore, while IR56d neurons that project posteriorly are responsive to both sucrose and HxA, those originating in taste pegs are selectively responsive to HxA, revealing two distinct functional classes of IR56d taste neurons.

**Fig 3.**
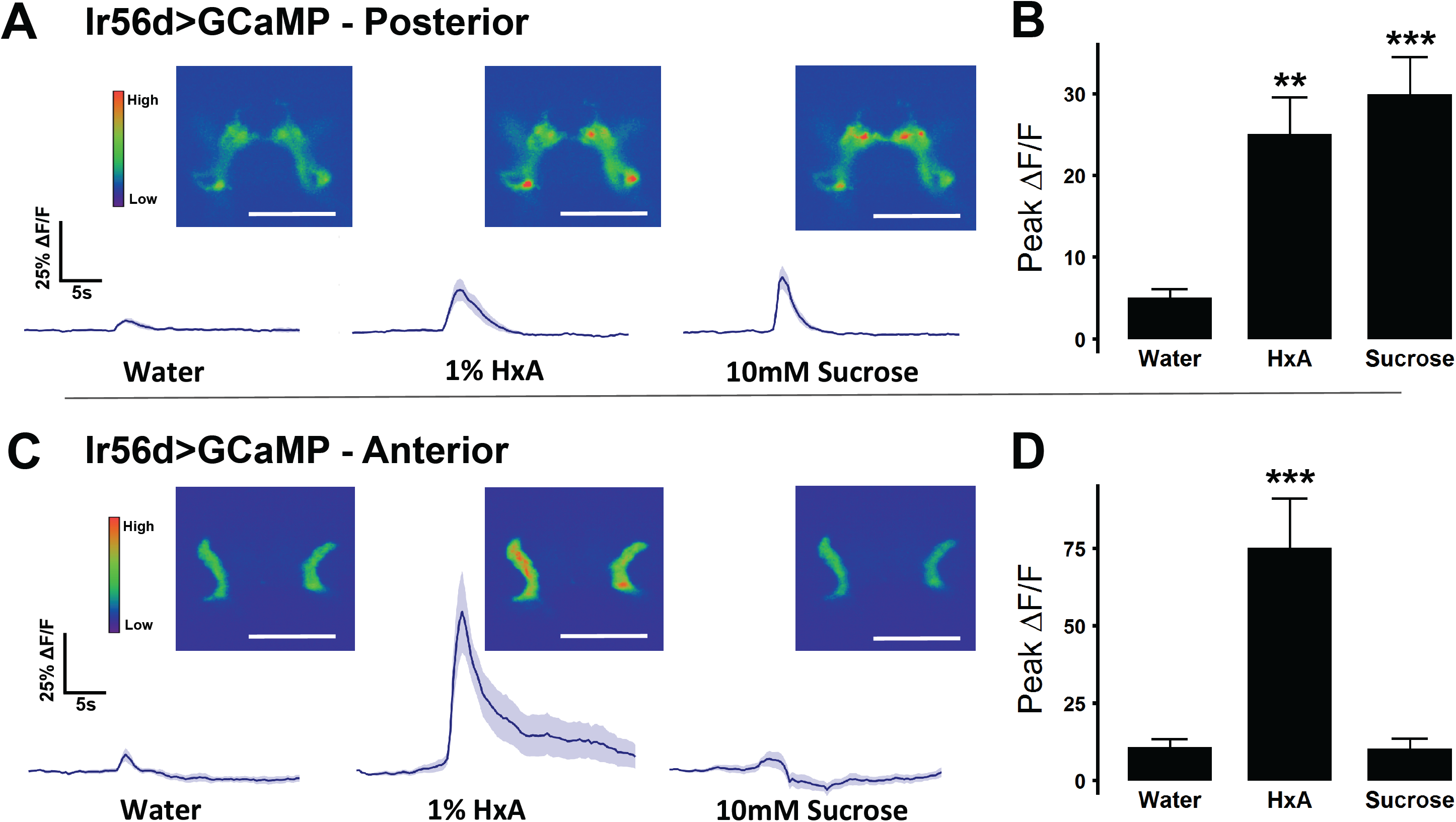
Response to sugar and fatty acid differs in anterior and posterior IR56d projections. **(A)** Activity traces and representative images of calcium activity in IR56d posterior projections in response to water, 1% HxA, and 10mM sucrose (n = 13 for each tastant). Shaded region of trace indicates +/- SEM **(B)** Average peak ΔF/F for data shown in (A). Error bars indicate SEM. One-way ANOVA with Tukey’s HSD - **p< 0.01, ***p< 0.001. **(C)** Activity traces and representative images of calcium activity in IR56d posterior projections in response to water, 1% HxA, and 10mM sucrose (n = 12 for each tastant). Shaded region of trace indicates +/- SEM **(D)** Average peak ΔF/F for data shown in (C). Error bars indicate SEM. One-way ANOVA with Tukey’s HSD - ***p< 0.001.

Flies exhibit feeding responses upon presentation of several FA classes [19]. It is possible that the IR56d neurons are broadly responsive to different classes of FAs. Alternatively, different classes of FAs may be sensed by distinct, or partially overlapping, populations of sensory neurons. To test the responsiveness of IR56d neurons to different classes of FAs, we first compared behavioral responses of IR56d-silenced flies to short-chain pentanoic acid (5-carbon), medium-chain octanoic acid (8 carbon), and long chain oleic acid (18-carbon). As compared to control IR56d>imp-TNT flies, IR56d-silenced flies exhibited weaker responses to octanoic acid, but retained robust PER to pentanoic acid (Fig 4A), suggesting that PER to short chain FAs is not dependent on IR56d neurons. Oleic acid did not elicit strong PER in either. We directly examined activity induced by different FAs in IR56d neurons with *in vivo* Ca2+ imaging in IR56d-GAL4>GCaMP5 flies. Octanoic acid robustly activated posterior IR56d projections, while both pentanoic and octanoic acid activated anterior IR56d projections (Fig 4B,C). Oleic acid, which did not induce PER, also did not activate IR56d projections in either posterior or anterior regions (Fig 4B,C). Together, these findings show that IR56d neurons respond to short and medium chain FAs, and further, that sub-populations of IR56d neurons that are separately represented in anterior and posterior locations in the SEZ have distinct FA response profiles.

**Fig 4.**
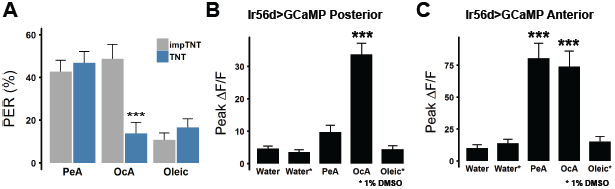
IR56d neurons are selectively responsive to short and medium chain FAs. **(A)** PER in response to short chain pentanoic acid (PeA, n = 46), medium chain octanoic acid (OcA, n = 26), and long-chain oleic acid (n = 46) in control imp-TNT flies. Blocking synaptic release in IR56d neurons with TNT significantly decreases PER to octanoic acid (n = 29), but does not affect PER for pentanoic (n = 42) or oleic acid (n = 42). Wilcoxon Rank-Sum Test - ***p<0.001. **(B)** Average peak ΔF/F for the posterior projections of IR56d neurons in response to water, 1% DMSO, pentanoic acid, octanoic acid, and oleic acid (n = 9, 8, 9, 9, 8, respectively), and **(C)** the anterior projections (n = 9, 7, 9, 9, 7, respectively). Error bars indicate SEM. One way ANOVA with Tukey’s HSD - ***p<0.001

Since silencing of either IR56d or Gr64f neurons abolishes PER to hexanoic and octanoic acid, one possibility is that neurons co-expressing both IR56d and Gr64f are required for medium-chain FA response (Fig 5A). To validate co-expression of IR56d and Gr64f, we used the LexA system to label Gr64f neurons (Gr64f-LexA>LexOp-CD8:GFP) and the GAL4 system to label IR56d neurons (IR56d-GAL4>UAS-RFP) (Fig 5A). Examining SEZ projections revealed co-localization within the posterior SEZ, with no co-localization detected in the anterior SEZ, suggesting the posterior IR56d neurons co-express Gr64f and IR56d. To determine whether the IR56d+/Gr64f+ co-expressing neurons are required for FA taste, we used intersectional strategies to selectively silenced anterior IR56d neurons of the taste pegs. Specifically, we repressed TNT expression in IR56d+/Gr64f+ using Gr64f-LexA to drive expression of the GAL80 repressor (IR56d-GAL4>UAS-TNT; Gr64f-LexA>LexAop-GAL80) (Fig 5B). Selectively silencing IR56d neurons that do not overlap with Gr64f did not affect PER to HxA, or as expected, to sugar, and no difference was observed between flies expressing imp-TNT and TNT, suggesting the taste peg neurons are dispensable for the reflexive feeding response to FAs (Fig 5B). Flies lacking Gr64f-LexA, but still harboring a copy of LexAop-GAL80 (IR56D-GAL4-TNT; LexAop-GAL80/+), showed reduced PER as expected (Fig S1).

**Fig 5.**
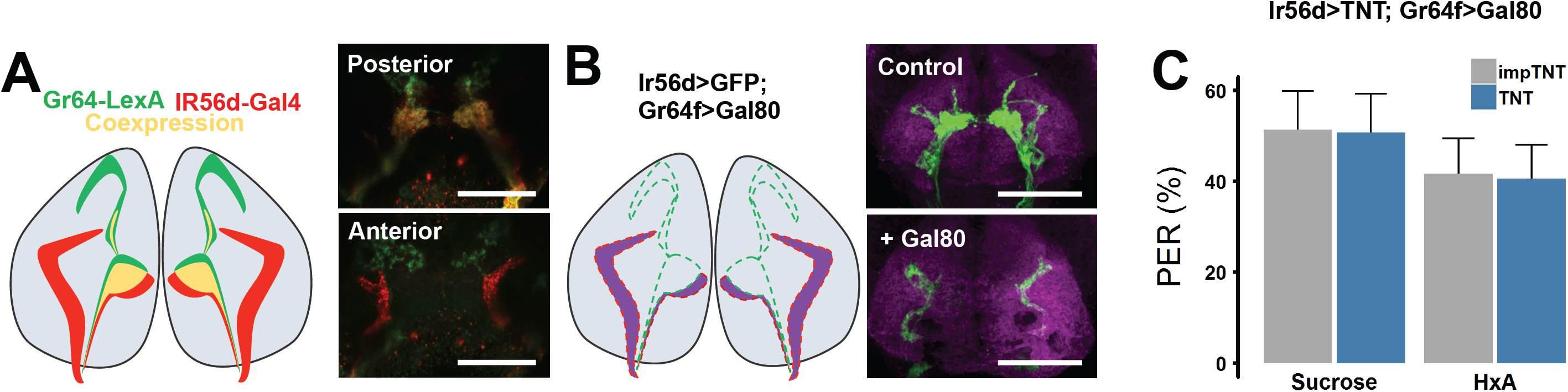
IR56d anterior projections are not required for PER to FAs. **(A)** IR56d and Gr64f neurons visualized with IR56d-GAL4 driving RFP and Gr64f-LexA driving GFP. Overlap can be seen in the posterior projections, but is minimal in anterior projections. **(B)** Driving the GAL4 repressor Gal80 with Gr64f-LexA limits GFP expression to the non-overlapping IR56d anterior projections. **(C)** Restricting TNT expression to the non-overlapping anterior projection neurons does not significantly impact PER to sugar or HxA (n = 23 imp-TNT; n = 22 TNT) Error bars indicate SEM. Wilcoxon Rank Sum Test (Sucrose: p> 0.98; HxA: p>0.96)

Although both sugars and FAs induce feeding behavior, it is unclear whether flies can qualitatively differentiate between these two classes of appetitive tastants. We have developed an assay in which an appetitive tastant is applied to the tarsi, paired with bitter quinine application to the proboscis, and the suppression of PER in subsequent response to the appetitive tastant is measured [40,45]. To determine whether flies can differentiate between sugars and FAs, we applied sucrose (conditioned stimulus) to the tarsi followed immediately with quinine application (unconditioned stimulus) to the proboscis. Following three training trials, memory was tested by application of sucrose (control) or HxA to the tarsi, in the absence of quinine, and measuring PER (Fig 6B). We also performed reciprocal experiments in which flies were trained with HxA and tested for PER to sucrose or HxA.

**Fig 6.**
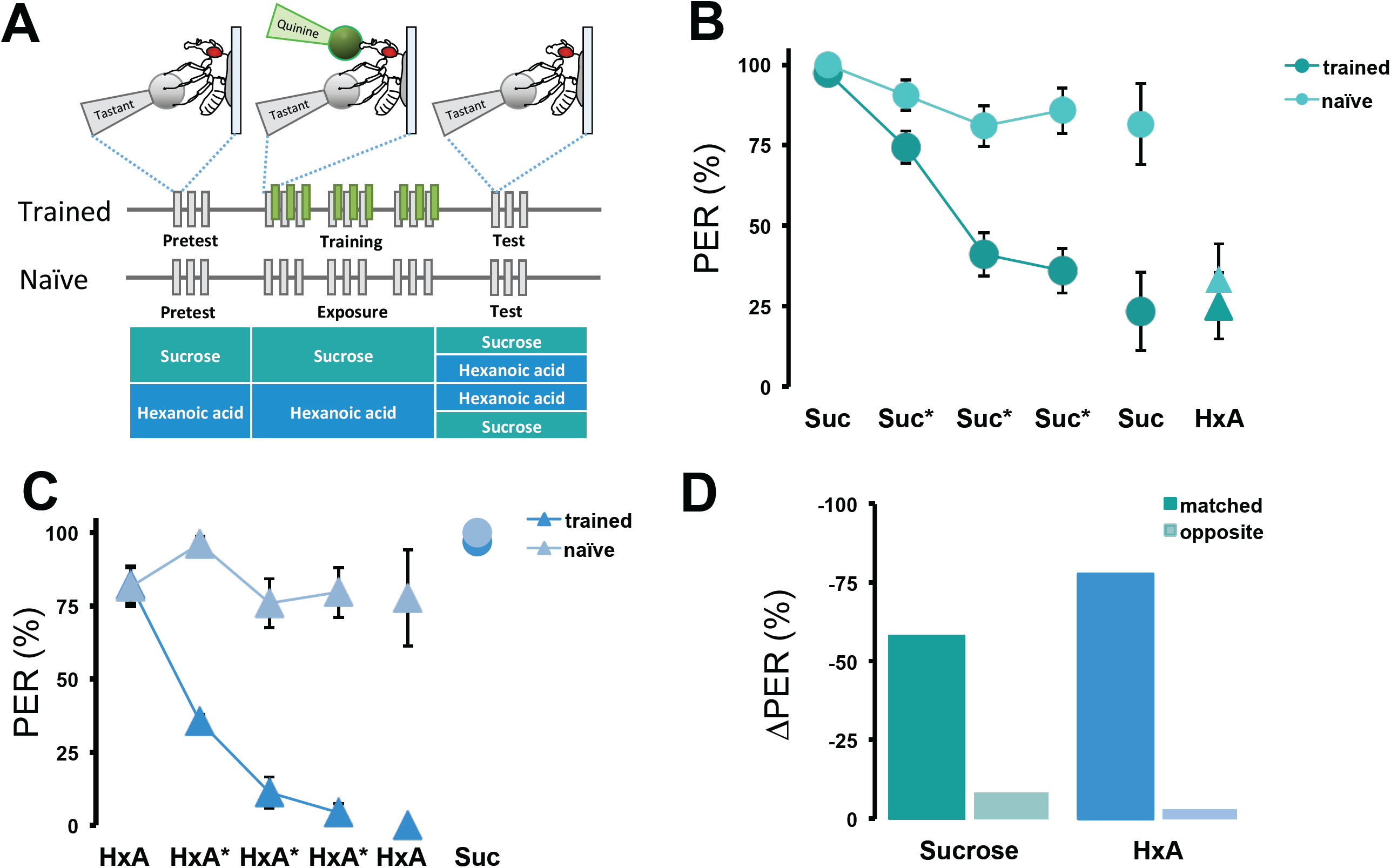
*Drosophila* discriminate between sucrose and HxA. **(A)** Taste memory protocol to determine sugar and FA discrimination. Flies are trained by pairing HxA or sucrose on tarsi with quinine on proboscis. PER in response to HxA and sucrose is then tested following training to sucrose or HxA in the absence of quinine. In control experiments, the same procedure is followed but quinine is never offered for measurement of naïve decline of tastant responses. **(B)** The pairing of sucrose and quinine (dark green circles) results in a significant reduction in PER across all three training trials compared to unpaired naïve flies provided sucrose alone (light green circles). PER response to sucrose in the test is significantly lower in trained flies compared to naïve flies (p=<0.001, n=7,11), but no differences in response to HxA (triangles) is detected between the experimental and naïve groups (p>0.31, n=11, 12). **(C)** The pairing of HxA and quinine (dark blue triangles) results in a significant reduction in PER across all three training trials compared to unpaired naïve flies provided HxA alone (light green triangles). The test PER response to HxA alone is significantly lower in trained flies compared to naïve flies (p<0.001, n=13,16), but no differences in response to sucrose (circles) is detected between the groups (p>0.59, n=12,12). **(D)** Comparing the percentage change in PER suppression reveals that flies train and tested to the same tastant (either sucrose or HxA) significantly reduced PER compared to flies trained and tested with different tastants.

As previously reported, pairing sugar with quinine significantly reduced PER over the course of three training trials and this memory persisted in the absence of quinine ([46,47]; Fig 6B). Further, there were no differences in PER to HxA between flies repeatedly provided sucrose in the absence of quinine and flies trained with sucrose-quinine pairing, indicating that the aversive taste memory formed to sucrose is not generalized to HxA. Conversely, the pairing of HxA and quinine resulted in a robust PER reduction that was not generalized to sucrose (Fig. 6C). Quantification of the percentage reduction in PER reveals that the ‘matched’ groups, where quinine is paired with an attractive tastant suppressed PER by 79-94%, while there was little PER suppression when the tested tastant was unmatched from the one previously paired with quinine (Fig 6D). This reciprocal discrimination between sucrose and HxA is different from the unilateral discrimination between two sugars reported previously [45] and indicates that flies can discriminate between sucrose and HxA based on their identity. Thus, sugars and FAs act as independent taste modalities in flies.

## Discussion

Here, we identify a small population of neurons within the taste system that is responsive to FAs. Other taste modalities, such as sweet and bitter, have been well studied in flies and mammals, but much less is known about FA taste. We previously identified a requirement for Gr64f-GAL4 neurons in the taste of FA [19]. The findings presented here further localize the reflexive feeding response to taste of hexanoic and octanoic acid, both medium-chain FAs, to a subpopulation of IR56d neurons. ‘Sweet-sensing’ neurons in *Drosophila* have been broadly classified as ones responding to sugars and other sweet tastants such as glycerol, subsets of which may also be involved in sensing other appetitive tastants such as amino acids [17,28,48]. Our findings indicate that IR56d neurons represent channels for FA detection, and likely allow flies to discriminate between different attractive tastants.

### Localization of fatty acid-sensitive neurons

We identify a subpopulation of sweet sensing neurons that are responsive to FAs. We have previously shown that genetic silencing or ablation of most sweet-sensing neurons using Gr64f-GAL4 abolished FA taste, suggesting shared neurons detect sugar and FA taste [19]. In flies, subpopulations of Gr64f neurons are selectively sensitive to different tastants including a Gr64e population that is sensitive to glycerol [48] and Gr5a subset that is sensitive to trehalose [17]. To localize the Gr64f neurons responsible for FA taste we conducted a targeted screen and silenced neurons that are believed to overlap with Gr64f neurons, which led us to study the IR56d population of neurons. ISilencing Ir56d neurons appears to selectively disrupt HxA taste without affecting response to sucrose, supporting the notion that independent channels within the Gr64f population mediate response to sugar and FAs. It is possible that FA-sensitive neurons are broadly tuned to FAs, or selectively respond to distinct classes of FAs. Our Ca2+ imaging experiments indicate that IR56d neurons are responsive to medium-chain HxA (six carbons (C6), saturated) and octanoic acid (C8, saturated) in both anterior and posterior regions, and to short-chain pentanoic acid (C5, saturated), but only in the anterior projections. We do not find IR56d neurons responsive to long chain, unsaturated oleic acid (C18, mono-unsaturated). These findings are supported by behavioral data revealing that flies exhibit robust PER in response to pentanoic acid, HxA, and octanoic acid, but not oleic acid. Therefore, it seems likely that flies are selectively responsive to short/medium chain FAs, but less responsive to long-chain and/or unsaturated FAs. It is possible that a population of taste neurons independent from IR56d are activated by pentanoic acid because it elicited a robust behavioral response that was independent of Ir56d neurons. Further, IR56d neurons may be activated by long-chain FAs that were not tested, and these could modulate feeding response and induce PER. Nevertheless, the findings presented here reveal specificity for short/medium chain FAs within a defined population of taste neurons, and further specificity for medium chain FAs in a subset of those neurons.

### Discrimination between attractive substances

The robust PER response induced by two different medium-chain FAs, hexanoic and octanoic acid, suggests they may be part of the fruit flies diet. Typical dietary fats, including many plant based oils, such as coconut oil, are rich in longer chain FAs including palmitic acid, oleic acid and linoleic acid [49]. However medium chain FAs are present in fermenting fruits such as guava and pollen, and it is possible this provides a substrate for *Drosophila* in the wild [50,51]. While less is known about the food substrates of *D. melanogaster,* the closely related *D. sechelia* consume Morinda fruit, which is rich in FAs [52,53]. Further, we have previously shown that a diet of HxA alone is sufficient for survival [19]. Therefore, it is possible that FA attraction evolved to promote consumption of calorically rich fruits consumed by *Drosophila.*

Our aversive memory assay indicates flies can discriminate between these attractive tastants. Sugar induces broad activation of Gr64f neurons, while the activation induced by HxA appears more restricted, and therefore it is possible that differences in activation allow for differentiation [54]. Alternatively, we find that HxA also activates IR56d neurons that emanate from the taste pegs and do not co-express Gr64f, so it is possible that the activation of these neurons by HxA allows discrimination. Finally, considering the different biochemical pathways involved in sugar and FA sensing [19], their identity can be coded by unique temporal and spatial dynamics of sensory neuron activation [16,55,56]. Differences in activation are suggested to provide a mechanism for olfactory discrimination within defined neural populations, and it is possible that similar mechanisms are utilized for attractive tastants [53]. In *Drosophila*, attractive tastants have been found to induce a wide range of excitatory responses ranging from acute to prolonged firing [29,57], providing a potential mechanism for discrimination. While the sensillar response to FAs has not been reported, the differences in Ca2+ response to sugar or HxA presentation within the SEZ suggest differences in temporal activation.

We find that flies can discriminate between sugar and FAs, but it is not known whether they can discriminate qualitatively between different classes of FAs. A previous study examining discrimination between different sugars found that flies are unable to discriminate based on quality, but could discriminate based on perceived palatability [45]. Here, we find that pentanoic acid elicits a robust PER response that is independent of IR56d neurons. The findings that distinct populations of neurons respond to FAs from different classes raises the possibility that flies may discriminate based on pattern of neuronal activation.

### Role of Phospholipase C in fatty acid taste

We have previously reported evidence that Phospholipase C (PLC) signaling in sweet-sensing Gr64f neurons is required for FA taste [19]. Flies with mutation or knockdown of the PLC-ß ortholog *no receptor potential A* (*norpA*) do not respond to HxA or octanoic acid but respond normally to sugars, indicating that distinct intercellular signaling mechanism underlie response to FAs and sugars [19,58]. Our findings that the IR56d-Gr64f co-expressing neurons respond to both sugars and FAs provide a system to directly test the role of *norpA* in neuronal response within a defined neuronal population. Further, our findings show that PER response to pentanoic acid does not require IR56d neurons, suggesting distinct populations of taste neurons regulate response to short-chain (pentanoic acid) and medium-chain FAs (HxA and octanoic acid). Testing the response of *norpA* deficient flies to short-chain FAs will inform whether PLC is more generally required for FA taste, or is only specific to medium-chain FAs detected by IR56d neurons.

### Higher order neurons involved in fatty acid perception

While much is known about taste coding within the SEZ, less is known about the higher order circuits governing taste. Sweet sensing neurons project to the antennal mechanosensory and motor center (AMMC) and downstream PAM dopaminergic neurons that are activated by sugar [39,59]. In addition, a separate populations of dopamine neurons, the PPL1 cluster, is required for olfactory appetitive memory and taste aversive conditioning [60–62]. A central question will be whether these higher order neurons downstream of the SEZ are also sensitive to FAs, or selectively tuned to sugar. To date, higher order neurons responsive to FA taste have not been identified. It is possible that sugar and FA taste signal through shared higher order dopaminergic neurons or, alternatively, each taste modality may activate distinct populations of higher order neurons that convey valence to the mushroom bodies, the memory and sensory integration center in *Drosophila* [63–65].

## Conclusions

Taken together, this study provides insight into the coding of FAs within the fly gustatory system. Our results reveal a population of sweet-sensing neurons that are tuned for medium-chain FAs, but not short- or long-chain FAs. Further, the finding that flies are capable of discriminating between FAs and sugars suggests coding differences, either at the level of spatial or temporal neuronal activation, that provides a mechanism to distinguish between tastants of the same valence. Understanding how FAs are coded within the fly brain provides a model for understanding taste in more complex systems and will offer insight into generalizable mechanisms for taste discrimination.

## Methods

### *Drosophila* maintenance and fly stocks

Flies were grown and maintained on standard food (New Horizon Jazz Mix, Fisher Scientific). Flies were maintained in incubators (Powers Scientific; Dros52) at 25°C on a 12:12 LD cycle, with humidity set at 55-65%. The background control line used in this study is the *w*^1118^ fly strain unless otherwise noted. The following fly straits were ordered from The Bloomington Stock Center, UAS-imp-TNT (28840), UAS-TNT (28838), UAS-TRPA1 (26263); UAS-CaMPARI; (58762); UAS-GFP (32186); UAS-GCaMP5 (42037); +;; Gr43a-GAL4 (57637), Gr5a-GAL4 (57591), Gr61a-GAL4 (57658), Gr64a-GAL4 (57662), Gr64c-GAL4 (57663), Gr64d-GAL4 (57665), Gr64e-GAL4 (57666), Gr64f-GAL4 (57668), IR56b-GAL4 (60706), IR56d-GAL4 (60708), LexAop-Gal80 (32213). Gr64f-LexA was kind gift from H. Tanimoto and previously described in [20]. Seven to nine day old mated female flies were used for all experiments in this study, except when noted.

### Proboscis Extension Reflex

For the initial screen using TNT, and specific testing of tarsal response, PER was measured by applying tastant to the tarsi, as previously described [19]. For all other PER experiments, including validation of *Ir56d* phenotypes, tastant was applied to the proboscis to match behavioral results with functional imaging. Flies were anesthetized on CO_2_, and mounted in a pipette tip so that their head and proboscis, but not tarsi, were exposed. After a 30-60m acclimation period, flies were presented with water and allowed to drink freely until water satiated. Flies that did not stop responding to water within 5 minutes were discarded. A small Kimwipe (Kimberley Clark) wick saturated with tastant was manually applied to the tip of the proboscis for 1-2s and proboscis extension reflex was monitored. Only full extensions were counted as a positive response. Each tastant was presented three times, with 10 seconds between each trial. Between tastant trials, the proboscis was washed with water and flies were allowed to drink. PER response was calculated as a percentage of proboscis extensions to total number of tastant stimulations. For experiments examining the effects of TRPA1 activation on PER, flies were mounted on a microscope slide using nail polish as described previously [19]. Flies were then placed on a heat plate heated to 34 °C and video of activity was recorded for 1 minute. The number of flies for each genotype showing PER within the 1 minute trial period was counted and the percentage of flies showing PER calculated.

### *In vivo* functional imaging

Flies were anaesthetized on ice and placed in a small chamber with the head and proboscis accessible. A small hole was cut in tin foil and fixed to the stage leaving a window of cuticle exposed, then sealed using dental glue (Tetric Flow – Ivoclar Vivadent). The proboscis was extended and a small amount of dental glue was used to secure it in place, ensuring the same position throughout the experiment.

The cuticle and connective tissue was dissected to expose the SEZ, which was bathed in artificial hemolymph (140 mM NaCl, 2mM KCl, 4.5 mM MgCl2, 1.5mM CaCl2, and 5mM HEPES-NaOH with pH = 7.0). Mounted flies were placed under a confocal microscope (Nikon A1) and imaged using a 25x water-dipping objective lens. The pinhole was opened to allow a thicker optical section to be monitored. Recordings were taken at 4Hz with 256 resolution. Tastants were delivered to the proboscis for 1-2s with a KimWipe wick operated by microcontroller (brand, type). For analysis, regions of interest (ROI) were drawn manually, taking care to capture the same area between control and experimental. Baseline fluorescence was recorded over 5 frames, 10s prior to tastant application. ΔF/F was calculated for each frame as (fluorescence – baseline fluorescence)/baseline fluorescence * 100. Average fluorescence traces were created by taking the average and standard error of ΔF/F for all samples per frame.

### Immunohistochemistry

Fly brains were dissected in ice-cold PBS and fixed in 4% formaldehyde, PBS,0.5% Triton-X 100 for 30 minutes. Brains were rinsed 3X with PBS, Triton-X for 10 minutes and incubated overnight at 4°C in NC82 (Iowa Hybridoma Bank [66]). The brains were rinsed again in PBS-TritonX, 3X for 10 minutes and placed in secondary antibodies (Donkey anti-Mouse 555; Life Technologies) for 90 minutes at room temperature. The brains were mounted in Vectashield (VectorLabs) and imaged on confocal microscope. Brains were imaged in 2um sections and are presented as the Z-stack projection through either the entire brain, or anterior and posterior regions of IR56d projections in the SEZ.

### Aversive Taste Memory

PER induction was performed in one-week old mated females as described previously [5, 16]. One-week-old mated females were collected and placed on fresh food for 24 h, then starved for 48 h in food-vials on wet Kimwipe paper. Flies were later anaesthetized on CO_2_ pad and glued using nail polish (Cat#72180, Electron Microscopy Science) by their thorax and wing base on a microscopy slide and left to recover in a box with wet Kimwipe paper for 3-6 h prior to experiments. For experiments, the slide was mounted vertically under the dissecting microscope (Olympus SZX12) and PER was observed. Flies were satiated with water before and during the experiment. Flies that did not initially satiate within 5 minutes were excluded from conditioning. We used a 1ml syringe (Tuberculine, Becton Dickinson & Comp) for tastant presentation. We used purified water, 10 mM and 1000 mM sucrose, 0.4% hexanoic acid and 10 mM quinine hydrochloride solutions. The protocol was adapted from [40].Here, for pretest, each fly was given 10 mM sucrose or 0.4% HxA on their tarsi three times with 10 sec inter-trial interval and the number of full proboscis extensions was recorded. During training, same stimulation as before was followed by a droplet of 10 mM quinine was placed on extended proboscis and flies were allowed to drink it for up to 2 sec or until they retracted their proboscis. After each session, tarsi and proboscis were washed with water and flies were allowed to drink to satiation. After training, flies were tested with either that same substance without quinine or with the untrained substance (matched or opposite trained groups). Another group of flies was measured as described above but quinine was never presented (naïve groups). At the end of each experiment, flies were given 1000mM sucrose to check for retained ability of PER and non-responders were excluded [11].

### Reagents

Sucrose was purchased from Fisher Scientific (FS S5-500). All other tastants were purchased from Sigma Aldrich. Sucrose, hexanoic acid (SA 153745), octanoic acid (SA C2875), pentanoic Acid (SA 240370) and quinine hydrochloride (SA 145904) were diluted in H_2_0. Oleic Acid (SA O1008) was diluted in DMSO 1% DMSO (Sigma).

### Statistical Analysis

All statistical tests were performed in R. For normally distributed data, Student’s t-test or ANOVA with Tukey’s post-hoc comparison was performed. For data that did not fit a normal distribution, nonparametric tests were used: Wilcoxon Rank-Sum or Kruskal-Wallis with Dunn’s post-hoc. Fisher’s Exact Test was used for binary categorical data. For all tests with multiple comparisons, Bonferroni p-value adjustment was performed.

## Acknowledgements

We are grateful to Michael Gordon (University of British Columbia), Seth Tomchik (Scripps, Florida), Anupama Dahanukar (University of California, Riverside) and Hiromu Tanimoto (Tohoku University, Japan) for technical support. This work was supported by NIH Grants 1R01 NS085252 to ACK and NSF IOS-1426265 to PM and ACK.

## Supplemental Data

**Table S1. A neuronal silencing screen for taste neurons sensitive to HxA**

**(A)** Data for PER in response to 100mM sucrose in flies with silenced populations of gustatory neurons by driving expression of TNT with the indicated. The ‘normalized’ column represents the PER value for experimental fly divided by the result for w1118. **(B)** Data for PER in response to 1% HxA. p-values computed by Kruskal-Wallis followed by Dunn’s post-hoc with Bonferroni p-value adjustment, with comparison to w1118 control flies. *p<0.05, **p<0.01, ***p<0.001.

**Fig S1. Flies continue to suppress PER in response to FAs in absence of Gr64f to drive LexAop-Gal80**

**(A)** PER for flies expressing either imp-TNT or TNT in IR56d neurons, (n = 31 for both groups), Without Gr64f-LexA to drive the LexAop-GAL80, as in Fig 5c, TNT is expressed in all IR56d neurons and PER to HxA is suppressed. Wilcoxon Rank Sum Test ***p<0.001

## References

1. Tepper BJ, Nurse RJ. Fat perception is related to PROP taster status. Physiol Behav. 1997;61: 949–954. doi:10.1016/S0031-9384(96)00608-7

2. Ramirez I. Chemosensory similarities among oils: does viscosity play a role? Chem Senses. 1994;19: 155–68. Available: http://www.ncbi.nlm.nih.gov/pubmed/8055265

3. Kinney NE, Antill RW. Role of olfaction in the formation of preference for high-fat foods in mice. Physiol Behav. 1996;59: 475–478. doi:10.1016/0031-9384(95)02086-1

4. Greenberg D, Smith GP. The controls of fat intake. Psychosom Med. 1996;58: 559–69. Available: http://www.ncbi.nlm.nih.gov/pubmed/8948004

5. Takeda M, Sawano S, Imaizumi M, Fushiki T. Preference for corn oil in olfactory-blocked mice in the conditioned place preference test and the two-bottle choice test. Life Sci. 2001;69: 847–854. doi:10.1016/S0024-3205(01)01180-8

6. Fukuwatari T, Shibata K, Iguchi K, Saeki T, Iwata A, Tani K, et al. Role of gustation in the recognition of oleate and triolein in anosmic rats. Physiol Behav. 2003;78: 579–583. doi:10.1016/S0031-9384(03)00037-4

7. Hiraoka T, Fukuwatari T, Imaizumi M, Fushiki T. Effects of oral stimulation with fats on the cephalic phase of pancreatic enzyme secretion in esophagostomized rats. Physiol Behav. 2003;79: 713–717. doi:10.1016/S0031-9384(03)00201-4

8. Han T, Lean M. No A clinical perspective of obesity, metabolic syndrome and cardiovascular disease. JRSM Cardiovasc Dis. 2016;25: 2048.

9. Rada P, Avena NM, Barson JR, Hoebel BG, Leibowitz SF. A High-Fat Meal, or Intraperitoneal Administration of a Fat Emulsion, Increases Extracellular Dopamine in the Nucleus Accumbens. Brain Sci. 2012;2: 242–253. doi:10.3390/brainsci2020242

10. Vosshall LB, Stocker RF. Molecular architecture of smell and taste in Drosophila. Annu Rev Neurosci. 2007;30: 505–533. doi:10.1146/annurev.neuro.30.051606.094306

11. Hallem EA, Dahanukar A, Carlson JR. Insect odor and taste receptors. Annu Rev Entomol. 2006;51: 113–135. doi:10.1146/annurev.ento.51.051705.113646

12. Liman ER, Zhang Y V., Montell C. Peripheral coding of taste. Neuron. 2014. pp. 984–1000. doi:10.1016/j.neuron.2014.02.022

13. Zhang Y, Hoon MA, Chandrashekar J, Mueller KL, Cook B, Wu D, et al. Coding of sweet, bitter, and umami tastes: Different receptor cells sharing similar signaling pathways. Cell. 2003;112: 293–301. doi:10.1016/S0092-8674(03)00071-0

14. Yarmolinsky D a., Zuker CS, Ryba NJP. Common Sense about Taste: From Mammals to Insects. Cell. 2009;139: 234–244. doi:10.1016/j.cell.2009.10.001

15. Masek P, Keene AC. Drosophila Fatty Acid Taste Signals through the PLC Pathway in Sugar-Sensing Neurons. PLoS Genet. 2013;9: e1003710. doi:10.1371/journal.pgen.1003710

16. Freeman EG, Dahanukar A. Molecular neurobiology of Drosophila taste. Curr Opin Neurobiol. 2015;34: 140–148. doi:10.1016/j.conb.2015.06.001

17. Marella S, Fischler W, Kong P, Asgarian S, Rueckert E, Scott K. Imaging taste responses in the fly brain reveals a functional map of taste category and behavior. Neuron. 2006;49: 285–295. doi:10.1016/j.neuron.2005.11.037

18. Thorne N, Chromey C, Bray S, Amrein H. Taste Perception and Coding in Drosophila. Curr Biol. 2004;14: 1065–1079. doi:10.1016/j.cub.2004.05.019

19. Masek P, Keene AC. Drosophila fatty acid taste signals through the PLC pathway in sugar-sensing neurons. PLoS Genet. 2013;9: e1003710. doi:10.1371/journal.pgen.1003710

20. Thoma V, Knapek S, Arai S, Hartl M, Kohsaka H, Sirigrivatanawong P, et al. Functional dissociation in sweet taste receptor neurons between and within taste organs of Drosophila. Nat Commun. Nature Publishing Group; 2016;7: 10678. doi:10.1038/ncomms10678

21. Jiao Y, Moon SJ, Wang X, Ren Q, Montell C. Gr64f Is Required in Combination with Other Gustatory Receptors for Sugar Detection in Drosophila. Curr Biol. 2008;18: 1797–1801. doi:10.1016/j.cub.2008.10.009

22. Flood TF, Iguchi S, Gorczyca M, White B, Ito K, Yoshihara M. A single pair of interneurons commands the Drosophila feeding motor program. Nature. 2013;499: 83–7. doi:10.1038/nature12208

23. Marella S, Mann K, Scott K. Dopaminergic Modulation of Sucrose Acceptance Behavior in Drosophila. Neuron. 2012;73: 941–950. doi:10.1016/j.neuron.2011.12.032

24. Pool AH, Kvello P, Mann K, Cheung SK, Gordon MD, Wang L, et al. Four GABAergic interneurons impose feeding restraint in Drosophila. Neuron. 2014;83: 164–177. doi:10.1016/j.neuron.2014.05.006

25. Wang Z, Singhvi A, Kong P, Scott K. Taste representations in the Drosophila brain. Cell. 2004;117: 981–991. doi:10.1016/j.cell.2004.06.011

26. Hussain A, Zhang M, Üçpunar HK, Svensson T, Quillery E, Gompel N, et al. Ionotropic Chemosensory Receptors Mediate the Taste and Smell of Polyamines. PLoS Biol. 2016;14: e1002454. doi:10.1371/journal.pbio.1002454

27. Fischler W, Kong P, Marella S, Scott K. The detection of carbonation by the Drosophila gustatory system. Nature. 2007;448: 1054–1057. doi:10.1038/nature06101

28. Zhang Y V., Ni J, Montell C. The Molecular Basis for Attractive Salt-Taste Coding in Drosophila. Science (80-). 2013;340: 1334–1338. doi:10.1126/science.1234133

29. Charlu S, Wisotsky Z, Medina A, Dahanukar A. Acid sensing by sweet and bitter taste neurons in Drosophila melanogaster. Nat Commun. 2013;18: 1199–1216. doi:10.1016/j.micinf.2011.07.011.Innate

30. Fujii S, Yavuz A, Slone J, Jagge C, Song X, Amrein H. Drosophila sugar receptors in sweet taste perception, olfaction, and internal nutrient sensing. Curr Biol. 2015;25: 621–627. doi:10.1016/j.cub.2014.12.058

31. Miyamoto T, Chen Y, Slone J, Amrein H. Identification of a Drosophila Glucose Receptor Using Ca2+ Imaging of Single Chemosensory Neurons. PLoS One. 2013;8: 1–8. doi:10.1371/journal.pone.0056304

32. Weiss LA, Dahanukar A, Kwon JY, Banerjee D, Carlson JR. The molecular and cellular basis of bitter taste in Drosophila. Neuron. 2011;69: 258–272. doi:10.1016/j.neuron.2011.01.001

33. Akerboom J, Chen T-W, Wardill TJ, Tian L, Marvin JS, Mutlu S, et al. Optimization of a GCaMP Calcium Indicator for Neural Activity Imaging. J Neurosci. 2012;32: 13819–13840. doi:10.1523/JNEUROSCI.2601-12.2012

34. Akerboom J, Chen T-W, Wardill TJ, Tian L, Marvin JS, Mutlu S, et al. Optimization of a GCaMP Calcium Indicator for Neural Activity Imaging. J Neurosci. 2012; doi:10.1523/JNEUROSCI.2601-12.2012

35. Sweeney ST, Broadie K, Keane J, Niemann H, O’kane CJ. Targeted Expression of Tetanus Toxin Light Chain in Drosophila Specifically Eliminates Synaptic Transmission and Causes Behavioral Defects. Neuron. 1995;14: 341–351.

36. Koh TW, He Z, Gorur-Shandilya S, Menuz K, Larter NK, Stewart S, et al. The Drosophila IR20a Clade of Ionotropic Receptors Are Candidate Taste and Pheromone Receptors. Neuron. 2014;83: 850–865. doi:10.1016/j.neuron.2014.07.012

37. Ganguly A, Pang L, Duong VK, Lee A, Schoniger H, Varady E, et al. A Molecular and Cellular Context-Dependent Role for Ir76b in Detection of Amino Acid Taste. Cell Rep. 2017;18: 737–750. doi:10.1016/j.celrep.2016.12.071

38. Croset V, Schleyer M, Gerber B, Benton R. Molecular and neuronal basis for amino acid sensing in the Drosophila larva. Sci Rep. 2016;6: 34871.

39. Kain P, Dahanukar A. Secondary Taste Neurons that Convey Sweet Taste and Starvation in the Drosophila Brain. Neuron. 2015;85: 819–832.doi:10.1016/j.neuron.2015.01.005

40. Keene AC, Masek P. Optogenetic induction of aversive taste memory. Neuroscience. 2012. pp. 173–180. doi:10.1016/j.neuroscience.2012.07.028

41. Hamada FN, Rosenzweig M, Kang K, Pulver SR, Ghezzi A, Jegla TJ, et al. An internal thermal sensor controlling temperature preference in Drosophila. Nature. 2008;454: 217–220. doi:10.1038/nature07001

42. Pulver SR, Pashkovski SL, Hornstein NJ, Garrity P a, Griffith LC. Temporal dynamics of neuronal activation by Channelrhodopsin-2 and TRPA1 determine behavioral output in Drosophila larvae. J Neurophysiol. 2009;101: 3075–3088. doi:10.1152/jn.00071.2009

43. Zhang W, Ge W, Wang Z. A toolbox for light control of Drosophila behaviors through Channelrhodopsin 2-mediated photoactivation of targeted neurons. Eur J Neurosci. 2007;26: 2405–2416. doi:10.1111/j.1460-9568.2007.05862.x

44. Inagaki HK, Jung Y, Hoopfer ED, Wong AM, Mishra N, Lin JY, et al. Optogenetic control of Drosophila using a red-shifted channelrhodopsin reveals experience-dependent influences on courtship. Nat Methods. 2013;11: 325–332. doi:10.1038/nmeth.2765

45. Masek P, Scott K. Limited taste discrimination in Drosophila. Proc Natl Acad Sci U S A. 2010;107: 14833–14838. doi:10.1073/pnas.1009318107

46. Masek P, Worden K, Aso Y, Rubin G, Keene A. A dopamine-modulated neural circuit regulating aversive taste memory in Drosophila. Curr Biol. 2015;25: 1535– 41.

47. Kirkhart C, Scott K. Gustatory learning and processing in the Drosophila. J Neurosci. 2015;15: 5950–8.

48. Wisotsky Z, Medina A, Freeman E, Dahanukar A. Evolutionary differences in food preference rely on Gr64e, a receptor for glycerol. Nat Neurosci. 2011;14: 1534– 1541. doi:10.1038/nn.2944

49. Orsavova J, Misurcova L, Vavra Ambrozova J, Vicha R, Mlcek J. Fatty acids composition of vegetable oils and its contribution to dietary energy intake and dependence of cardiovascular mortality on dietary intake of fatty acids. Int J Mol Sci. 2015;16: 12871–12890. doi:10.3390/ijms160612871

50. Bhat R, Suryanarayana LC, Chandrashekara KA, Krishnan P, Kush A, Ravikumar P. Lactobacillus plantarum mediated fermentation of Psidium guajava L. fruit extract. J Biosci Bioeng. 2015;119: 430–432. doi:10.1016/j.jbiosc.2014.09.00.

51. Kaplan M, Karaglu O, Eroglu N, Silici S. Fatty Acid and Proximate Composition of Bee Bread. Food Technol Biotechnol. 2016;54: 497–504.

52. Wang M, Kikuzaki H, Csiszar K, Boyd CD, Maunakea A, Fong SFT, et al. Novel trisaccharide fatty acid ester identified from the fruits of Morinda citrifolia (noni). J Agric Food Chem. 1999;47: 4880–4882. doi:10.1021/jf990608v

53. Kim HK, Kwon MK, Kim JN, Kim CK, Lee YJ, Shin HJ, et al. Identification of novel fatty acid glucosides from the tropical fruit Morinda citrifolia L. Phytochem Lett. 2010;3: 238–241. doi:10.1016/j.phytol.2010.09.002

54. Chu B, Chui V, Mann K, Gordon MD. Presynaptic gain control drives sweet and bitter taste integration in Drosophila. Curr Biol. 2014;24: 1978–1984. doi:10.1016/j.cub.2014.07.020

55. Reiter S, Campillo Rodriguez C, Sun K, Stopfer M. Spatiotemporal Coding of Individual Chemicals by the Gustatory System. J Neurosci. 2015;35: 12309– 12321. doi:10.1523/JNEUROSCI.3802-14.2015

56. Carleton A, Accolla R, Simon SA. Coding in the mammalian gustatory system. Trends in Neurosciences. 2010. pp. 326–334. doi:10.1016/j.tins.2010.04.002

57. Ling F, Dahanukar A, Weiss L a, Kwon JY, Carlson JR. The molecular and cellular basis of taste coding in the legs of Drosophila. J Neurosci. 2014;34: 7148– 64. doi:10.1523/JNEUROSCI.0649-14.2014

58. Hardie RC, Martin F, Chyb S, Raghu P. Rescue of Light Responses in the Drosophila " Null " Phospholipase C Mutant, norpA P24, by the Diacylglycerol Kinase Mutant, rdgA, and by Metabolic Inhibition*. J Biol Chem. 2003;278: 18851–18858. doi:10.1074/jbc.M300310200

59. Liu C, Plaçais P-Y, Yamagata N, Pfeiffer BD, Aso Y, Friedrich AB, et al. A subset of dopamine neurons signals reward for odour memory in Drosophila. Nature. Nature Publishing Group; 2012;488: 512–516. doi:10.1038/nature11304

60. Das G, Lin S, Waddell S. Remembering Components of Food in Drosophila. Front Integr Neurosci. 2016;10: 4. doi:10.3389/fnint.2016.00004

61. Scaplen KM, Kaun KR. Reward from bugs to bipeds: a comparative approach to understanding how reward circuits function. J Neurogenet. 2016;30: 133–48. doi:10.1080/01677063.2016.1180385

62. Masek P, Keene A. Gustatory processing and taste memory in Drosophila. J Neurogenet. 2016;30: 112–21.

63. Aso Y, Siwanowicz I, Bräcker L, Ito K, Kitamoto T, Tanimoto H. Specific dopaminergic neurons for the formation of labile aversive memory. Curr Biol. 2010;20: 1445–1451. doi:10.1016/j.cub.2010.06.048

64. Claridge-Chang A, Roorda RD, Vrontou E, Sjulson L, Li H, Hirsh J, et al. Writing Memories with Light-Addressable Reinforcement Circuitry. Cell. 2009;139: 405– 415. doi:10.1016/j.cell.2009.08.034

65. Lewis LPC, Siju KP, Aso Y, Friedrich AB, Bulteel AJB, Rubin GM, et al. A Higher Brain Circuit for Immediate Integration of Conflicting Sensory Information in Drosophila. Curr Biol. 2015;25: 2203–2214. doi:10.1016/j.cub.2015.07.015

66. Wagh DA, Rasse TM, Asan E, Hofbauer A, Schwenkert I, Dürrbeck H, et al. Bruchpilot, a protein with homology to ELKS/CAST, is required for structural integrity and function of synaptic active zones in Drosophila. Neuron. 2006;49: 833–844. doi:10.1016/j.neuron.2006.02.008

